# Pharmacologically diverse antidepressant drugs disrupt the interaction of BDNF receptor TRKB and the endocytic adaptor AP-2

**DOI:** 10.1101/591909

**Authors:** Senem Merve Fred, Liina Laukkanen, Cecilia A Brunello, Liisa Vesa, Helka Goos, Iseline Cardon, Rafael Moliner, Tanja Maritzen, Markku Varjosalo, Plinio C Casarotto, Eero Castrén

## Abstract

Antidepressant drugs activate TRKB (tropomyosin-related kinase B), however it remains unclear whether these compounds employ a common mechanism for achieving this effect. We found by using mass spectrometry that the interaction of several proteins with TRKB was disrupted in the hippocampus of fluoxetine-treated animals (single intraperitoneal injection), including members of the AP-2 complex (adaptor protein complex-2) involved in vesicular endocytosis. The interaction of TRKB with the cargo-docking mu subunit of the AP-2 complex (AP2M) was disrupted by both acute and repeated fluoxetine treatment. However, while the coupling between full length TRKB and AP2M was disrupted by fluoxetine, the interaction between AP2M and the TRKB C-terminal peptide was resistant to this drug, indicating that the binding site targeted by fluoxetine must lie outside of the TRKB:AP2M interface. In addition to fluoxetine, other pharmacologically diverse antidepressants imipramine, rolipram, phenelzine, ketamine, and the ketamine metabolite 2R,6R-hydroxynorketamine (RR-HNK) also decreased the interaction between TRKB:AP2M *in vitro*, as measured by ELISA. Silencing the expression of AP2M in MG87.TRKB cell line led to increased surface positioning of TRKB and to a higher response to BDNF (brain-derived neurotrophic factor), observed as the levels of active TRKB. Moreover, animals haploinsufficient for the *Ap2m1* gene displayed increased levels of active TRKB *in vivo*, as well as an enhanced cell surface expression of the receptor in cultured hippocampal neurons.

Taken together, our data suggests that disruption of the TRKB:AP2M interaction is an effect shared by several antidepressants with diverse chemical structures and canonical modes of action.

## Introduction

Compromised plasticity has been suggested as one of the major causes of depression (1). Supporting this idea, morphological and functional deficits have been observed in the brains of patients with mood disorders (2–4). Reduced neurotrophic support, such as by BDNF (brain-derived neurotrophic factor) signaling via TRKB, has been linked to the atrophy observed in these patients in the neuronal networks regulating mood and cognition (5, 6).

Antidepressant drugs (AD) have been suggested to act by improving neuronal connectivity, plasticity, and information processing mainly by targeting monoamine neurotransmission, such as serotonin (5-HT) and noradrenaline (4). In line with this, an intact BDNF-TRKB signaling system is crucial for the efficacy of antidepressant compounds, as mice overexpressing a dominant negative TRKB isoform (TRKB.T1) or haploinsufficient for BDNF (*Bdnf+/-*) are not responsive to AD (7). Furthermore, the activation of TRKB receptor signaling, and putatively the reinstatement of plasticity, is associated with the response to AD (7–9).

The binding of BDNF to TRKB initiates dimerization and phosphorylation of tyrosine residues in the intracellular portion of the receptor (10) which allows docking of adaptor proteins to these sites. Phosphorylated tyrosine 515, for example, serves as a docking site for SHC adaptor protein, while phospholipase C gamma (PLC-γ1) binds to the phosphorylated Y816 residue (11). Interestingly, the phosphorylation of TRKB receptors at Y706/7 and Y816 residues are enhanced by several antidepressants in adult mouse hippocampus while no change was detected at the Y515 residue (7, 9). Despite the evidence on the necessity of the BDNF/TRKB system for antidepressant responses, the mechanism by which these drugs might trigger TRKB activation remains unclear.

TRKB is mainly localized intracellularly in vesicles that are translocated to neuronal surface through neuronal activity (12–14). In this scenario, proteins able to modulate TRKB transit to and from the cell surface, therefore modulating its exposure to BDNF, posed as an interesting possibility to understand the antidepressant-induced effects. In line with this idea, our mass spectrometry-based screen for TRKB interactors that change upon fluoxetine treatment identified the AP-2 complex as TRKB interactor. AP-2 is crucial for clathrin-dependent endocytosis, and participates in the removal of several transmembrane receptors, and their ligands from the cell surface (15). AP-2 is a heterotetrameric complex which consists of two large chains (alpha- and beta-adaptin), one medium chain (mu, AP2M), and one small chain (sigma) (16). TRK receptors have been identified among the targets for rapid clathrin-mediated endocytosis upon binding to their ligands (17). Earlier studies demonstrated that BDNF-induced phosphorylation of TRKB receptors recruits AP-2 complex and clathrin to the plasma membrane in cultured hippocampal neurons (18). More recently, a novel relationship has been established between TRKB receptors and AP-2, suggesting that autophagosomes containing activated TRKB are transported to the cell soma in an AP-2-dependent manner (19).

In the present study, we identified a putative binding site on the TRKB receptor for AP2M, which provides a known interface for interaction between these proteins. Thus, we aimed at investigating whether the interface between TRKB and AP2M is a site of action for AD. Specifically, we tested the effects of several AD such as fluoxetine, imipramine, rolipram, phenelzine, ketamine, and RR-HNK as well as non-antidepressants with psychoactive effects (diazepam and chlorpromazine), on the interaction of TRKB receptors with AP2M.

## Methods

### Animals

Adult (7 weeks old) male C57BL/6RccHsd mice from Harlan (Netherlands), kept with free access to food and water, were used for TRKB immunoprecipitation and mass spectrometry analysis. Briefly, the animals received a single intraperitoneal (i.p.) injection of fluoxetine (30 mg/kg) and were euthanized by CO_2_. An independent cohort of female and another one of male mice from the same supplier (12-18 weeks old) received fluoxetine for 7 days or single i.p. injection, respectively. The hippocampi were collected and processed for enzyme-linked immunosorbent assay (ELISA) and co-immunoprecipitation (co-IP) as described below. All procedures were in accordance with international guidelines for animal experimentation and the County Administrative Board of Southern Finland (ESLH-2007-09085/Ym-23, ESAVI/7551/04.10.07/2013).

The generation of floxed AP2M mice carrying loxP sites in front of exon 2 and behind exon 3 of the AP2M encoding gene *Ap2m1* is described in literature (20). These conditional knockout (KO) mice were crossed with CMV-Cre deleter mice to achieve ubiquitous excision of exons 2 and 3. The resulting AP2M.het animals were crossed with AP2M.wt mice, thus excluding the generation of AP2M homozygous KO which are embryonically lethal (21). All animals were genotyped prior to experiments using PCR on genomic DNA extracted from tissue biopsies, and confirmed by Western blot analysis of hippocampus. The KO allele was detected using the forward primer GCTCTAAAGGTTATGCCTGGTGG and the reverse primer CCAAGGGACCTACAGGACTTC that generate a fragment of 404 bp (PCR at 58°C, 25 cycles). All experiments involving AP2M.het mice were reviewed and approved by the ethics committee of the “Landesamt für Gesundheit und Soziales” (LAGeSo) Berlin and were conducted according to the committee’s guidelines. For brain dissection, animals (15 weeks old) were anesthetized with isoflurane prior to cervical dislocation. The hippocampus was dissected on ice, and processed as described below.

### Drugs

Acetylsalicylic acid (aspirin, #A2093, Sigma-Aldrich), fluoxetine (#H6995, Bosche Scientific), imipramine (#I7379-5G, Sigma-Aldrich), phenelzine (#P6777, Sigma-Aldrich), rolipram (#R6520, Sigma-Aldrich), ketamine (#3131, Tocris), 2R,6R-hydroxynorketamine (#6094, Tocris), diazepam (#D0899, Sigma-Aldrich), chlorpromazine (#C8138, Sigma-Aldrich), recombinant human BDNF (#450-02, Peprotech), and k252a (TRK inhibitor, #K2015, Sigma-Aldrich) were used (22, 23). All compounds were dissolved in DMSO for *in vitro* experiments, except BDNF, diluted in PBS. For *in vivo* administration, fluoxetine was diluted in sterile saline for intraperitoneal single injection (7) or in the drinking water for repeated administration regimen (8).

### Immunoprecipitation and mass spectrometry

For TRKB proteome quantitative mass spectrometry analysis, adult mouse hippocampal samples (n=3) were mechanically homogenized in NP lysis buffer [20 mM Tris-HCl; 137 mM NaCl; 10% glycerol; 50 mM NaF; 1% NP-40; 0.05 mM Na_3_VO_4_; containing a cocktail of protease and phosphatase inhibitors (#P2714 and #P0044, respectively, Sigma-Aldrich)]. The samples were centrifuged (16000 xg) at 4°C for 15 min, and the supernatant was subjected to pre-clearing (control) and TRKB immunoprecipitation using Protein-G sepharose containing spin columns (#69725, Pierce Spin Columns Snap Cap). Briefly, 30 μl of Protein-G sepharose were added to the columns and washed twice with 200 μl NP buffer. Samples (400 μg of total protein) were cleared through the columns without antibody (1 h, under rotation at 4°C). Unbound samples were collected (1000 xg, 1 min at 4°C) pre-incubated with 5 μl TRKB antibody (Goat anti-mouse TRKB Antibody, #AF1494, R&D Systems) and cleared through new Protein-G sepharose tubes. After 1 h rotation in the cold, the columns were washed twice with NP buffer and twice with TBS buffer (1000 xg, 1 min at 4°C). The proteins were then eluted from the column with 0.1 M glycine (3x 1000 xg, 1 min, 4°C), or with Laemmli buffer for samples submitted to Western blot; and stored at −80°C until further analysis. Cysteine bonds of the proteins were reduced with TCEP [Tris(2-carboxyethyl)-phosphine hydrochloride salt, #C4706 Sigma-Aldrich, USA] and alkylated with iodoacetamide (#57670 Fluka, Sigma-Aldrich, USA). Samples were digested with trypsin (Sequencing Grade Modified Trypsin, #V5111, Promega), and digested peptides were purified with C18 microspin columns (Harvard Apparatus, USA) (24).

Mass spectrometry analysis was performed on an Orbitrap Elite hybrid mass spectrometer (Thermo Fisher Scientific) coupled to EASY-nLC II system (Xcalibur version 2.7.0 SP1, Thermo Fisher Scientific). The peptides were separated with a 60 min linear gradient from 5 to 35% of buffer B (98% acetonitrile and 0.1% formic acid in MS grade water). The analysis was performed in data-dependent acquisition: a high resolution (60 000) FTMS full scan (m/z 300-1700) was followed by top20 CID-MS2 scans in ion trap (Energy 35). The maximum fill time was 200 ms for both FTMS (Full AGC target 1000000) and the ion trap (MSn AGC target of 50000). The MS/MS spectra were searched against the mouse component of the UniProtKB-database (release 2012_08, X entries) using SEQUEST search engine in Proteome Discoverer™ software (version 1.4, Thermo Fisher Scientific). Carbamidomethylation (+57.021464 Da) of cysteine residues was used as static modification, and oxidation (+15.994491 Da) of methionine was used as dynamic modification. Precursor mass tolerance and fragment mass tolerance were set to <15 ppm and <0.8 Da, respectively. A maximum of two missed cleavages was allowed, and results were filtered to a maximum false discovery rate (FDR) of 0.05. The final hit table was rendered based on the following criteria: the average signal from three controls from Sepharose columns only ≤0.33, specific signal minus signal from Sepharose column ≥2.

### Cell culture and silencing of AP2M expression

The cell line 3T3 (mouse fibroblasts) stably transfected to express TRKB (MG87.TRKB) were used for *in vitro* assays (23). MG87.TRKA cell line was used for antibody validation. The cells were maintained at 5% CO2, 37°C in Dulbecco’s Modified Eagle’s Medium [DMEM, containing 10% fetal calf serum, 1% penicillin/streptomycin, 1% L-glutamine, and 400 μg/ml G418 (only for MG87.TRKB)].

For silencing AP2M expression, MG87.TRKB cells were transfected with a mix of 4 AP2M specific shRNA sequences (#TG712191, Origene, USA or scrambled sequence as control) using Lipofectamine 2000 (#11668019, Thermo Fisher Scientific) as transfecting agent according to the manufacturer’s instructions. The expression of GFP reporter was confirmed by microscopy 24 h after transfection, and AP2M expression levels were determined by Western blot. The cells were challenged with BDNF (0 or 25 ng/ml/15 min) and lysed for ELISA or Western blot as described below.

Primary cultures of cortical and hippocampal cells from rat embryos (E18) or P0 mice (for AP2M.het and wt) were prepared as described in the literature (25), and maintained in NeuroBasal medium (Thermo Fisher Scientific) supplemented with 1% B-27 supplement (Thermo Fisher Scientific), 1% penicillin/streptomycin, and 1% L-glutamine at 37°C (25).

### Sample processing and Western blot

Samples from hippocampi of male and female mice were sonicated in NP lysis buffer and centrifuged (16000 xg at 4°C for 15 min). The supernatants were either submitted to TRKB:AP2M ELISA, or processed for immunoprecipitation of TRKB or AP2M, and submitted to SDS-PAGE. Supernatant of samples from AP2M.wt and AP2M.het mice, as well as samples from MG87.TRKB cells were also submitted to SDS-PAGE. For cell-free ELISA assays described below, the samples from MG87.TRKB cells were acidified with 1 M HCl (pH ∼2) at room temperature (RT) for 15 min, and the pH was restored to 7.4 with 1 M NaOH. This procedure aims to disrupt non-covalent interactions between proteins in a given complex.

The resolved proteins were transferred to polyvinylidene difluoride (PVDF) membranes, blocked with 3% BSA (bovine serum albumin) in TBST [20 mM Tris-HCl; 150 mM NaCl; 0.1% Tween-20; pH 7.6] for 2 h at RT and incubated overnight (ON) at 4°C with antibodies against TRKB (1:1000, #AF1494, R&D Systems); AP2M (1:1000, #sc-515920, Santa Cruz), pTRKB.Y515 (1:1000, #9141, Cell Signaling), pTRKB.Y706/7 (1:1000, #4621, Cell Signaling), pTRKB.Y816 (1:1000, #4168, Cell Signaling) or GAPDH (1:5000, #2118, Cell Signaling or #ab8245, Abcam). The membranes were then incubated with HRP-conjugated secondary antibodies (1:5000, anti-Gt, #61-1620, Invitrogen; anti-Rb, #170-5046, Bio-Rad; or mouse IgG kappa binding protein, #516102, Santa Cruz), and the HRP activity was detected by incubation with ECL and exposure to a CCD camera. The signal from each target was subtracted from background, normalized by GAPDH or TRKB (145 kDa) and expressed as percentage of AP2M.wt, scrambled or vehicle-treated groups.

### AP2M:TRKB interaction and surface TRKB ELISA assays

MG87.TRKB cells were incubated with fluoxetine, imipramine, rolipram, aspirin, ketamine, RR-HNK (1-100 μM), phenelzine (0.01-100 μM), chlorpromazine (10 μM) or diazepam (10 μM). An independent group received k252a (10 μM), followed 10 min later by fluoxetine (10 μM) or BDNF (25 ng/ml). The cells were lysed 15 min after the last drug administration. For comparative reasons, primary cultures of cortical and hippocampal cells from rat embryo (8-10 DIV) were incubated with fluoxetine (1-10 μM/15 min). The concentration ranges were chosen based on literature (23). Finally, mice received fluoxetine at 15 mg/kg for 7 days in drinking water (8) or at 30 mg/kg as a single i.p. injection 30 min (9) before sample collection, and the TRKB:AP2M interaction was analyzed by a sandwich ELISA method and/or co-IP using hippocampal lysates.

The AP2M:TRKB interaction was determined by ELISA based on the general method described in literature (23, 26), with minor adjustments to detect TRK:AP2M interaction. Briefly, white 96-well plates (OptiPlate 96F-HB, Perkin Elmer) were coated with capturing anti-TRK antibody (1:1000, #92991S, Cell Signaling or #AF1494, R&D Systems) in carbonate buffer (pH= 9.8) ON at 4°C. Following a blocking step with 3 % BSA in PBST [137 mM NaCl; 10 mM Phosphate; 2.7 mM KCl; 0.1% Tween-20; pH 7.4] (2 h, RT), samples were incubated ON at 4°C. The incubation with antibody against AP2M (1:2000, #sc-515920, Santa Cruz; ON, 4°C) was followed by HRP-conjugated mouse IgG kappa binding protein (1:5000, #516102, Santa Cruz; 2 h, RT). Finally, the chemiluminescent signal generated by the reaction with ECL was analyzed in a plate reader (Varioskan Flash, Thermo Fisher Scientific).

The surface levels of TRKB were also determined by ELISA (27) in MG87.TRKB cells treated with fluoxetine or imipramine (10 μM; 15 min or 2 h) and also after AP2M downregulation. Hippocampal neurons (10 DIV) of haploinsufficient *Ap2m* mice were also tested for their TRKB surface expression. Briefly, cells cultivated in clear bottom 96-well plates (ViewPlate 96, Perkin Elmer) were washed with ice-cold PBS and fixed with 100 μl of 4% paraformaldehyde (PFA) per well. After washing with PBS and blocking with PBS containing 5% non-fat dry milk and 5% BSA, the samples were incubated with primary anti-TRKB antibody (1:1000 in blocking buffer, #AF1494, R&D Systems) ON at 4°C. Following a brief washing with PBS, the samples were incubated with HRP-conjugated anti-Gt antibody (1:5000 in blocking buffer, #61-1620, Invitrogen) for 1 h at RT. The cells were washed 4x with 200 μl of PBS for 10 min each. Finally, the chemiluminescent signal generated by reaction with ECL was acquired in a plate reader.

The cell-free assays were performed in white 96-well plates. The plates were pre-coated with anti TRKB antibody (1:1000, #AF1494, R&D Systems) in carbonate buffer (pH 9.8), ON at 4°C. Following blocking with 3% BSA in PBST buffer (2 h at RT), 120 μg of total protein from each sample (treated as for cell-free ELISA) were added and incubated ON at 4°C under agitation. The plates were then washed 3x with PBST buffer, and fluoxetine (0.01-100 μM) was added for 15 min. Following the washing protocol, the secondary antibody against AP2M was added (1:2000, #sc-515920, Santa Cruz) ON at 4°C under agitation. The plates were then incubated with HRP-conjugated mouse IgG kappa binding protein (1:5000, #516102, Santa Cruz) for 2 h at RT. The HRP activity was detected by luminescence following incubation with ECL on a plate reader.

For the cell-free AP2M:TRKB C-terminal interaction, the plates were pre-coated with antibody against AP2M (1:500, #516102, Santa Cruz) in carbonate buffer, then samples (treated as for cell-free ELISA) from MG87.TRKB cells were added ON at 4°C. N-biotinylated synthetic peptides (ctrl, pY, or Y816A; Genscript, USA, sequences described in figure 1h) of the last 26 amino acids (aa) of rat TRKB (0.1 μg/ml= 34 nM, in PBST) were added for 1 h at RT; followed by HRP-conjugated streptavidin (1:10000, #21126, Thermo Fisher Scientific) for 1 h at RT. The luminescence was determined upon incubation with ECL as described. In an independent experiment, immobilized AP2M from MG87.TRKB lysates was incubated with pY TRKB C-terminal peptide and fluoxetine (0 or 10 μM) for 1 h at RT; and the luminescence was determined via HRP-conjugated streptavidin activity reaction with ECL by a plate reader.

**Figure 1.**
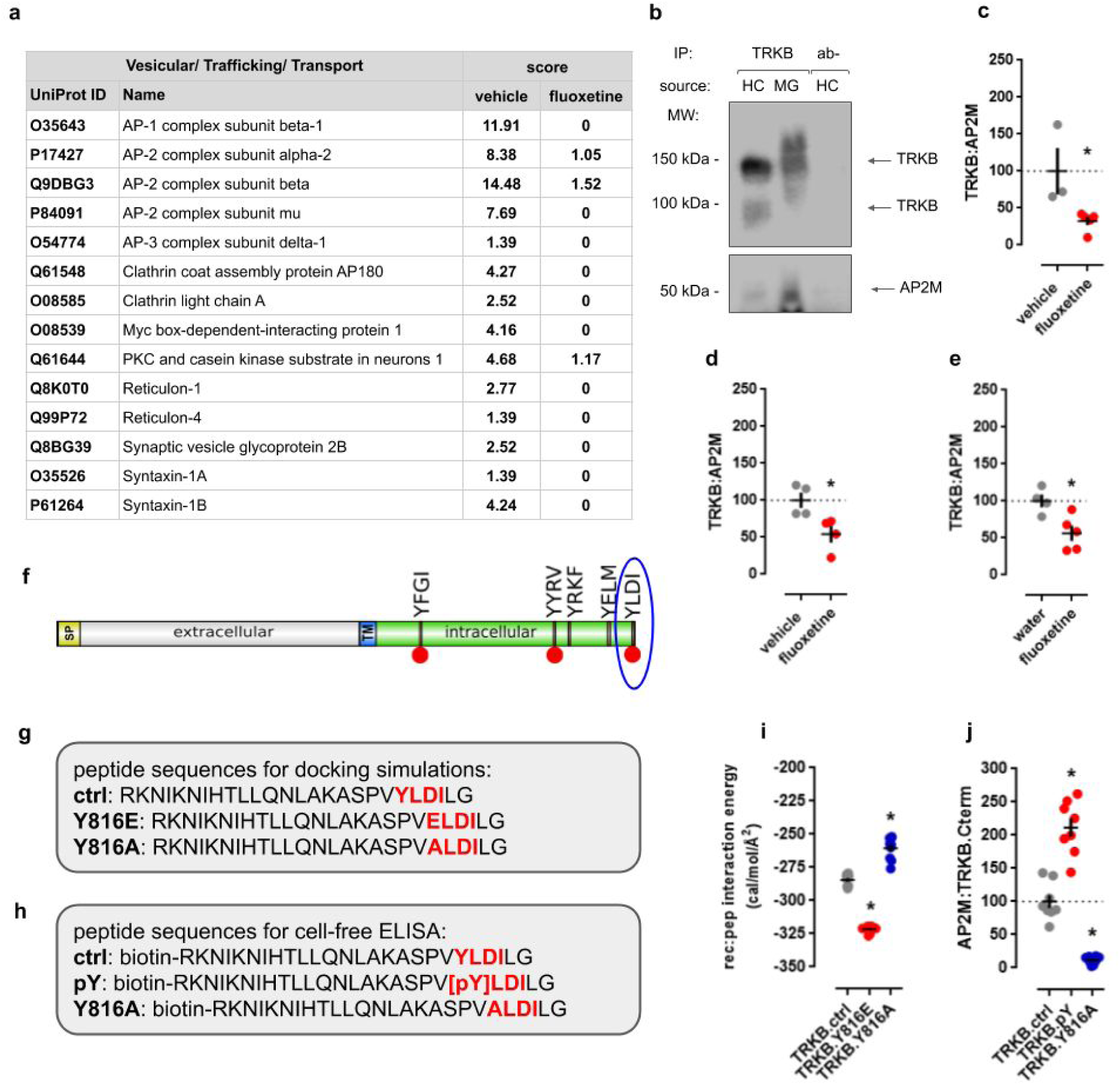
Interaction between TRKB and AP-2. **(a)** List of TRKB interactors involved in vesicle trafficking with mass spectrometry scores in vehicle- and fluoxetine-treated groups. For a full list of interacting proteins and the effects of fluoxetine administration, please refer to supplement tables 1 and 2 in the FigShare repository, respectively. **(b)** Co-IP of TRKB (145 or 100 kDa) and AP2M (50 kDa; IP:TRKB, WB:AP2M, TRKB) in samples from MG87.TRKB cells and hippocampus of 7 weeks old male mouse. Acute fluoxetine administration (30 min) decreases the TRKB:AP2M interaction in mouse hippocampus measured by **(c)** Western blot (IP:AP2M, WB:TRKB) or **(d)** ELISA, respectively. **(e)** Repeated administration of fluoxetine (7 days) also disrupts the TRKB:AP2M interaction, as measured by ELISA. **(f)** The C-terminal region of TRKB exhibits AP2M-binding motifs identified by Eukaryotic Linear Motif library (TRG_ENDOCYTIC_2): Y(515)FGI, Y(705)YRV, Y(726)RKF, Y(782)ELM, Y(816)LDI (blue circle). Yellow: SP, signal peptide; grey: extracellular portion (N-terminal); blue: TM, transmembrane; green: intracellular (C-terminal), red circles indicate the sites of Y-phosphorylation in TRKB. **(g)** The docking simulations were performed using sequences of the peptides spanning the last 26 amino acids of TRKB receptor. **(h)** Synthetic peptides (0.1 μg/ml= 34 μM) were used in cell-free ELISA to measure their binding to AP2M protein in MG.87.TRKB cell lysate. Red residues represent the TRG_ENDOCYTIC_2-positive motif and mutated sites. **(i)** The phosphomimetic peptide (TRKB.Y816E, red circles) shows lower free energy for the receptor:peptide docking simulations than wild-type peptide (TRKB.ctrl, grey circles), while TRKB.Y816A mutant (blue circles) displayed the highest free energy values (n=10/group). **(j)** Phosphorylated peptide (TRKB.pY, red circles) showed the higher interaction with AP2M compared to control peptide (TRKB.ctrl, grey circles), while TRKB.Y816A mutant (blue circles) displayed the lowest interaction levels as measured by cell-free ELISA (n=8/group). *p<0.05 from the wild-type peptide TRKB.ctrl. Black crosses represent mean/SEM.

The background-subtracted signal from each sample in each of the described assays was expressed as percentage of control group (vehicle-treated or WT peptide).

### Immunofluorescence staining and image analysis

The colocalization experiments of TRKB and the endosome markers RAB5, RAB7, and RAB11 (28) were performed in cultured cortical cells. Briefly, 15 DIV cells were fixed in 4% PFA (in PBS) after being treated with fluoxetine (1 μM, 15 min). The coverslips were washed in PBS and incubated in blocking buffer [5% normal donkey serum; 1% BSA 0.1% gelatin; 0.1% Triton X-100; 0.05% Tween-20 in PBS] for 1 h at RT. The cells were stained with anti-TRKB antibody (1:500, #AF1494, R&D Systems) and anti-RAB5 antibody (1:500, #NB120-13253, Novus Biologicals), anti-RAB7 antibody (1:500, #D95F2, Cell Signaling), or anti-RAB11A antibody (5 μg/mL, #71-5300, Thermo Fisher Scientific) ON at 4°C under agitation. Next, Alexa Fluor-conjugated secondary antibodies (Thermo Fisher Scientific) were diluted in PBS (1:1000). After a brief wash in PBS, the coverslips were incubated in donkey anti goat-647 for TRKB and donkey anti rabbit-568 for the RAB-specific antibodies for 45 min at RT and fixed in Dako Fluorescence Mounting Medium (S3023, Dako North America, Inc.). Control coverslips without primary antibody staining were also included.

Imaging was performed with a Leica TCS SP8 X inverted confocal microscope, 63x oil objective (1.40 NA) at 1024×1024 pixel resolution. At least 6 to 10 Z-stack steps were acquired with 0.30 μm intervals from each image. ImageJ FIJI Coloc2 plug-in (https://imagej.net/Coloc_2) was used for the colocalization analysis. The experimenters selected and analyzed the regions of interests (ROIs of spines or neurites) blindly to the treatments. The ROIs which passed the requirement of Costes p-value>0.95 were selected for further statistical analysis of Pearson’s correlation coefficient values from ctrl and fluoxetine groups.

The antibodies used in Western blot, ELISA or immunostaining were validated in previous studies (29–31), *e.g.* against pY residues of TRKB (7, 9, 23). The specificity of AP2M antibody was assessed in this study in transgenic mice and AP2M-silenced cells. For the antibody used to detect total TRKB we performed Western blot with samples from MG87 cells expressing TRKB or TRKA, and the results can be found on the FigShare repository.

### Structural bioinformatics data mining and docking assay

In order to address a putative interaction between AP2M and TRKB, we submitted the full length, canonical sequence of rat and mouse TRKB (Uniprot: Q63604 or P15209, respectively) to Eukaryotic Linear Motif library (32). Several TRG_ENDOCYTIC_2 motifs (TRG2; a tyrosine-based sorting signal responsible for the interaction with AP2M subunit) were found for both mouse and rat TRKB. Of particular interest, the Y816 residue is also activated by BDNF and antidepressants. Thus, the sequence of the last 26aa residues of rat TRKB (WT, Y/E or Y/A) and full length AP2M model generated through RaptorX server (33) were submitted to docking simulations using CABS-docking server (34).

### Statistical analysis

Data was analyzed by Student’s t test, one- or two-way ANOVA (or their non-parametric equivalents) followed by Fisher’s LSD *post hoc* test, when appropriate. Values of p<0.05 were considered significant. All data generated in the present study, including blot and immunostaining images, is stored in FigShare under CC-BY license, DOI:10.6084/m9.figshare.7976528.

## Results

### AP2M is an interaction partner of TRKB receptors

Samples from mouse hippocampus [vehicle (n=3) and fluoxetine-treated (30 mg/kg, single i.p. injection, n=3)] were processed for co-IP with TRKB-specific antibody and analyzed by mass spectrometry to investigate the TRKB interactome, and the changes in the interactome induced by acute fluoxetine administration. The complete list of interactors can be found in FigShare (supplement table 1). The list of identified interactors that are components of the synaptic and endocytic machineries are listed with their score from mass spectrometry in figure 1a. In the present study, we decided to focus on the interaction with AP-2 as a putative modulator of TRKB surface localization and activation since three out of four subunits of the complex were found associated with the TRKB receptor (figure 1a). Moreover, acute fluoxetine administration induced uncoupling of TRKB from AP-2 which is evidenced from the reduced scores of the AP-2 complex subunits alpha-2 (P17427), beta (Q9DBG3), and mu (P84091) in the samples from fluoxetine treated-mice compared to the vehicle-treated control animals (figure 1a).

Several cargo proteins docked to AP-2 complex are bound via recognition of single or multiple tyrosine-based motifs by the AP2M subunit. These motifs in the cytosolic tails of transmembrane receptors consist of four amino acid residues (35). The initial tyrosine residue of the motif is followed by two random amino acids, and the signal ends with one of the hydrophobic residues leucine, isoleucine, phenylalanine, valine or methionine: Y-X-X-φ (27). We searched for such tyrosine-based motifs in mature full length TRKB. The Eukaryotic Linear Motif library (32) indicated the presence of multiple putative tyrosine-based sorting signals [classified as TRG ENDOCYTIC 2 motifs - TRG2] in the cytosolic domain of the rat and mouse canonical TRKB sequence; Y(515)FGI, Y(705)YRV, Y(726)RKF, Y(782)ELM, Y(816)LDI as seen in figure 1f. Interestingly, one of the presumed AP2M binding sites, Y(816)LDI, which is located within the C-terminal portion of TRKB, is phosphorylated following antidepressant administration (9). Thus, we focused on the TRKB interaction with the cargo-docking mu subunit of the AP-2 complex (AP2M). We further confirmed the interaction between TRKB and AP2M in lysates of mouse hippocampus and the TRKB-overexpressing MG87 cell line by co-IP and Western Blot (figure 1b).

The acute administration of fluoxetine (30 mg/kg) decreased the TRKB:AP2M interaction, as measured by Western blot [t(6)= 2.805, p=0.0310, vehicle n=3, fluoxetine n=5] or ELISA [t(6)= 2.966, p=0.0251, vehicle n=4, fluoxetine n=4] (figure 1c,d). To find out whether the drug-induced disruption persists long-term with extended drug treatment, we examined the effect of 7 days exposure to fluoxetine in mice [in the drinking water (15 mg/kg)]. This regimen was chosen based on evidence indicating the effectiveness of fluoxetine in learned helplessness model in mice (36). Indeed, the decrease in the TRKB:AP2M coupling was also found in mouse hippocampus [t(7)=3.115; p=0.0170, vehicle n=4, fluoxetine n=5], as analyzed by ELISA (figure 1e). Furthermore, we validated the assay based on the decrease in the TRKB:AP2M interaction following the partial depletion of AP2M in MG87.TRKB cells [data and plot found in FigShare repository; mean/SD/n of percent from ctrl: 100.0/ 34.24/ 11; siAP2M: 47.40/ 25.62/ 10; t(19)= 3.952, p=0.0009].

In order to assess the interaction mechanism between AP2M and TRKB, we generated a model of full length AP2M in the RaptorX server (33, 37). For the docking simulations, the AP2M model and the last 26 aa residues from the C-terminal portion of TRKB (figure 1g), including variants with either a phospho-dead (Y816A) or a phosphomimetic mutation (Y816E), of the critical tyrosine were uploaded to CABS-dock server (34). We obtained the Gibbs’ free energy for the AP2M:TRKB peptide interaction, and the lowest 10 values from each group were used in statistical analysis. The interaction energy of the AP2M:TRKB.Y816E complex was lower than that for the AP2M:TRKB.ctrl complex, suggesting a more stable interaction, while it was significantly higher for the AP2M:TRKB.Y816A complex compared to AP2M:TRKB.ctrl [Kruskal-Wallis= 25.83; p<0.0001, n=10/group] (figure 1i). Based on these *in silico* results, we proceeded to analyze the AP2M:TRKB C-terminus interaction *in vitro* by ELISA. For this purpose, 3 variants of N-biotinylated synthetic peptides of the last 26 aa of rat TRKB were used (ctrl, pY or Y816A) (figure 1h). The interaction between AP2M and the phosphorylated TRKB peptide (TRKB.pY) was stronger compared to control peptide. In line with our *in silico* data, the AP2M:TRKB.Y816A interaction was weaker in comparison with the control peptide [Kruskal-Wallis= 20.48; p<0.0001, n=8/group] (figure 1j).

### Role of AP2M in TRKB function

Previous studies indicate that AP2M depletion in mammalian cells prevents the assembly of functional AP-2 complexes (38) and results in a loss of AP-2 function (17). Thus, we investigated the impact of AP2M downregulation on activation of TRKB receptor in MG87.TRKB cells overexpressing TRKB receptor. To induce TRKB signaling, the cells were challenged with BDNF following the silencing of AP2M, as there was no detectable phosphorylation of the receptor without BDNF. The depletion of AP2M [t(14)=5.521; p<0.0001, n=8/group] (figure 2a) was associated with a marginal decrease in TRKB levels [t(13)=1.901; p=0.0797, ctrl n=8, siAP2M n=7] (figure 2b). However, silencing of AP2M enhanced the effect of BDNF on TRKB phosphorylation at residues Y706/7 [t(14)=3.099; p=0.0078, n=8/group] (figure 2c) and Y816 [F(1,20)=12.21; p=0.0023, n=6/group] (figure 2e), but not at residue Y515 [t(8)=0.092; p=0.928, n=5/group] (figure 2d). The plasma membrane localization of TRKB receptors was also increased [t(28)=7.71; p<0.0001, n=15/group] (figure 2f) by the downregulation of AP2M in the absence of BDNF.

**Figure 2.**
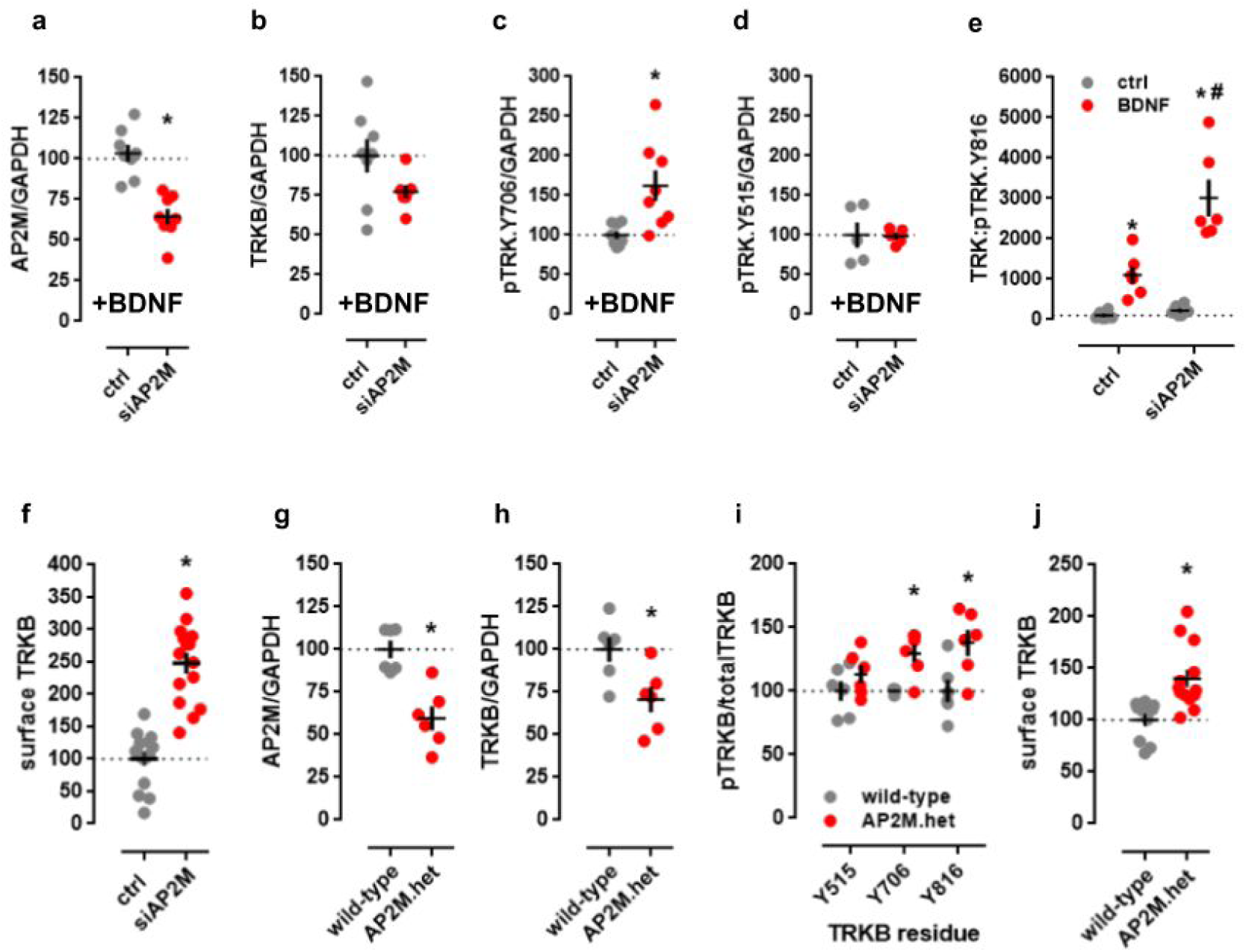
AP2M regulates TRKB cell surface localization and response to BDNF. Cultured MG87.TRKB cells were transfected with shRNA to silence the expression of AP2M and challenged with BDNF (0 or 25 ng/ml/15 min). Silencing of **(a)** AP2M (50 kDa) was associated with **(b)** decrease in total TRKB (145 kDa) levels and **(c)** increase in phosphorylation of TRKB at pY706 (145 kDa) but not in **(d)** pY515 (145 kDa) as measured by Western blot (n=5-8/group). **(e)** Phosphorylation of the pY816 site was augmented in siAP2M samples following the BDNF treatment as measured in ELISA (n=6/group). **(f)** The silencing of AP2M expression in MG87.TRKB led to an increased exposure of TRKB at the cell surface, as measured by surface ELISA (n=6/group). The Western blot analysis of protein samples from AP2M.het animals showed decreased levels of **(g)** AP2M and **(h)** total TRKB in the hippocampus, associated with **(i)** increased levels of phosphorylated TRKB at Y706 and Y816 residues (n=5-6/group). **(j)** Surface TRKB levels were also increased in primary cultures of hippocampal cells from AP2M.het mice in surface ELISA. *p<0.05 for comparison with ctrl/wild-type group; #p<0.05 for comparison with ctrl/siAP2M group. Black crosses: mean/SEM.

Consistently, the reduced level of AP2M in the hippocampus of haploinsufficient AP2M.het mice [t(10)=4.63; p=0.0009, n=6/group] (figure 2g) was associated with reduced levels of total TRKB in the hippocampus of these animals [t(10)=2.794; p=0.019, n=6/group] (figure 2h). However, we observed increased levels of pTRKB [genotype: F(1,29)=17.59; p=0.0002, n=6/group] at residues Y706/7 and Y816 (Fisher’s LSD, p=0.015 and 0.0016, respectively) (figure 2i). Similarly to the observed effects in MG87.TRKB cells, hippocampal cultures of AP2M.het animals (P0) showed increased levels of surface TRKB [t(21)= −3.603, p=0.002, WT n=10, AP2M.het n=13], figure 2j.

### Antidepressants and BDNF differentially affect the TRKB:AP2M interaction

The systemic administration of fluoxetine (30 mg/kg, i.p., 30 min) have a disruptive effect on the TRKB:AP2M interaction. We next tested whether other antidepressants similarly disrupt the TRKB:AP2M interaction. Indeed, the AD fluoxetine, imipramine, rolipram, phenelzine, ketamine, and RR-HNK reduced the TRKB:AP2M interaction in MG87.TRKB cells as measured by ELISA [fluoxetine: F(3,26)=4.554, p=0.011, n=12,6,6,6; imipramine: F(3,43)=7.883, p=0.0003, n=12,11,12,12; rolipram: F(3,19)=6.78, p=0.0027, n=6,6,6,5; phenelzine: Kruskal-Wallis=19.34, p=0.0017, n=18,12,12,12,6,6; ketamine: F(3,44)=7.421, p=0.0004, n=12,12,12,12; RR-HNK: F(3,35)= 4.022, p=0.0147, n=12,12,10,5], (Fisher’s LSD p<0.05), (figure 3a-f). However, the negative control aspirin did not induce a disruption [aspirin: F(3,40)=0.082, p=0.96, n=10,10,12,12] (figure 3g). Furthermore, the administration of non-AD with psychoactive profile such as chlorpromazine or diazepam failed to affect the TRKB:AP2M interaction [for chlorpromazine: t(42)=0.6143; p=0.5423, n=22,22; for diazepam: t(18)=1.167, p=0.2583, n=12,8] (figure 3h,i).

**Figure 3.**
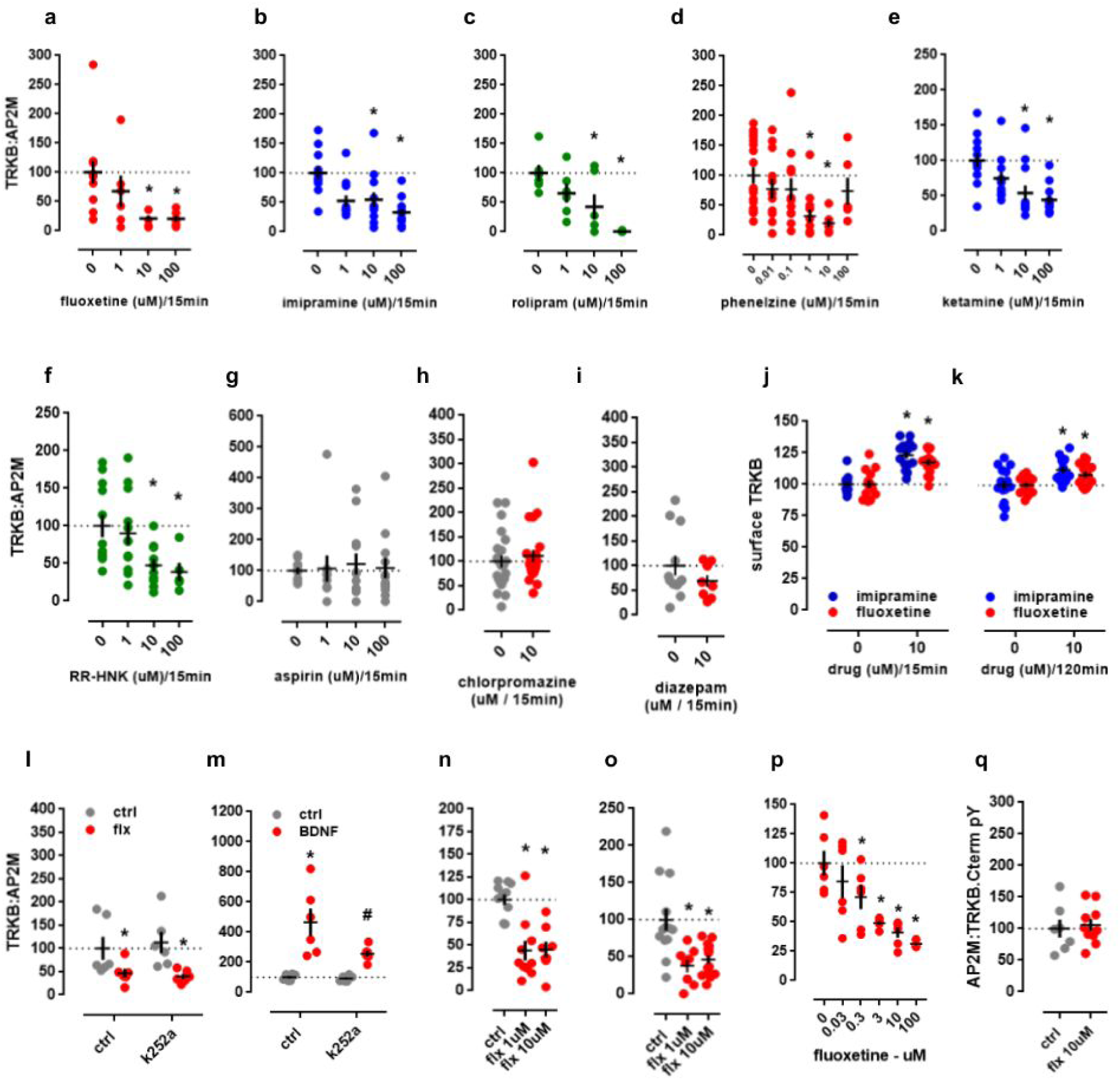
Antidepressant-induced disruption of the TRKB:AP2M interaction. MG87.TRKB cell cultures were treated with antidepressants **(a)** fluoxetine, **(b)** imipramine, **(c)** rolipram, **(d)** phenelzine, **(e)** ketamine or **(f)** RR-HNK, and the TRKB:AP2M interaction was determined by ELISA. The compounds **(g)** aspirin, **(h)** chlorpromazine or **(i)** diazepam were used as negative controls (n=6-22/group). **(j)** Surface levels of TRKB increased following fluoxetine or imipramine treatment as measured by surface ELISA (10 μM/15 min; n=15/group), and **(k)** surface levels of TRKB were still elevated 2 h after the administration of these drugs. ELISA measurements showed that the acute treatment of MG87.TRKB cells with k252a (10 μM) did not prevent **(l)** fluoxetine-(10 μM), but **(m)** BDNF-induced (25 ng/ml) effects on the TRKB:AP2M interaction (n=6-12/group). **(n)** Cortical and **(o)** hippocampal cells (8-10 DIV) were treated with fluoxetine (1-10 μM/15 min), and the levels of TRKB:AP2M determined by ELISA as described (n=8-13/group). As shown by a cell-free ELISA, fluoxetine was able to disrupt the AP2M interaction with **(p)** full length, but not with **(q)** TRKB C-terminal peptide in lysates from MG87.TRKB cells (n=6-10/group). *p<0.05 from ctrl/ctrl, water or 0uM, #p<0.05 from ctrl/k252a group. Black crosses: mean/SEM.

The administration of fluoxetine or imipramine increased the levels of surface exposed TRKB in MG87.TRKB cells as measured by ELISA [15 min: F(1,54)=62.54, p<0.0001, ctrl/fluoxetine n=14/14, ctrl/imipramine n=15/15; 120 min: F(1,56)=15.35, p=0.0002, n=15/group]. Interestingly, the disruptive effect of fluoxetine on the TRKB:AP2M interaction is not dependent on TRKB activation, since pre-treatment with the TRK inhibitor k252a was not able to prevent this effect [interaction effect: F(1,20)=0.3066, p=0.586, n=6/group] (figure 3l). BDNF administration (25 ng/ml) also resulted in an increased interaction of TRKB receptors with AP2M, but contrary to what was observed for antidepressants, this increase was dependent on TRKB activation, since pretreatment with k252a prevented this effect [interaction effect: F(1,20)=4.528, p=0.046, n=6/group] (figure 3m). Similarly to the observations for MG87.TRKB cells, fluoxetine was able to disrupt the AP2M:TRKB interaction in primary cultures of cortical [F(2,26)=14.75, p<0.0001, n=11,10,8] and hippocampal cells [F(2,32)=9.562, p=0.0006, n=13,8,13] of rat embryos (figure 3n,o).

Finally, we observed that fluoxetine disrupted the interaction between full length TRKB and AP2M in MG87.TRKB cell lysates as measured in a cell-free ELISA [F(5,30)=9.71; p<0.0001, n=6/group] with effective doses ranging from 0.3-100 μM (Fisher’s LSD, p<0.05 for all) (figure 3p). However, no effect of fluoxetine (10 μM) was observed on the interaction between AP2M and the TRKB C-terminal peptide [t(15)=0.35; p=0.73, ctrl n=7, flx n=10] (figure 3q) suggesting additional binding sites for fluoxetine outside of this region.

To the present point it is plausible to consider that disruption of TRKB:AP2M occurs in the cell surface as well as in endocytic vesicles. Therefore, to address the fate of TRKB receptors after an acute fluoxetine challenge, we determined the colocalization of TRKB with early, late and recycling endosome markers RAB5, RAB7 and RAB11, respectively (39, 40) (figure 4a,d,g). The Student’s t test indicates no significant effect of fluoxetine administration and the colocalization of TRKB and RAB proteins in neurites [RAB5: t(41)=0.8490, p=0.4008, ctrl n=26, fluoxetine n=17; RAB7: t(56)=0.7730, p=0.4428 ctrl n=30, fluoxetine n=28; RAB11: t(64)=0.8233, p=0.4134, ctrl n=37, fluoxetine n=29] of cortical neurons (figure 4c,f,i). No effect of fluoxetine treatment was observed in spines for TRKB:RAB5 [t(36)=0.5053, p=0.6164; ctrl n=19/group] (figure 4b) or TRKB:RAB11 [t(31)=1.561, p=0.1288, ctrl n=17, fluoxetine n=16] (figure 4h) colocalization. However, a significant effect of fluoxetine was observed on the colocalization of TRKB and RAB7 [t(36)=2.230, p=0.0321, ctrl n=19/group] (figure 4e).

**Figure 4.**
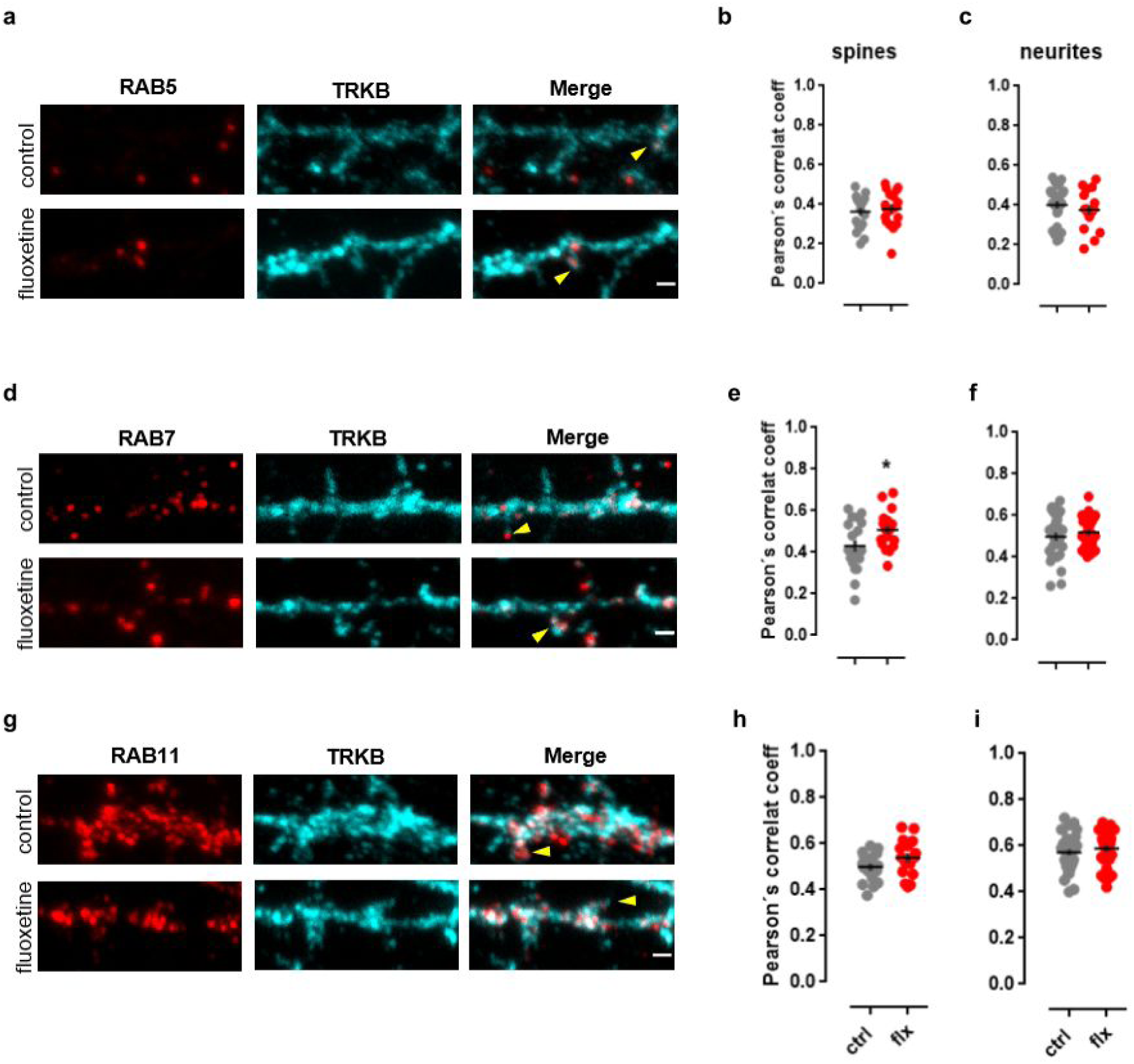
Effect of fluoxetine on TRKB colocalization with endosomal markers. The colocalization (indicated by the arrows) of TRKB with **(a-c)** early (RAB5), **(d-f)** late (RAB7) and **(g-i)** recycling (RAB11) endosome markers was determined in the spines and neurites of cultured cortical neurons (14-16 DIV) treated with fluoxetine (1 μM/15 min) (n=16-37 spines or neurite shafts /group). *p<0.05 from ctrl group. Black crosses indicate mean ± SEM. The scale bars indicate 1 μm.

## Discussion

Sub-chronic (41) and chronic fluoxetine treatment (42, 43) can induce changes in the proteomics profile in rodent brains. This compound altered the expression of proteins involved in neuroprotection, serotonin biosynthesis and axonal transport in hippocampus and frontal cortex (42), as well as in endocytosis and transport in visual cortex (43).

The present study identified the endocytic complex AP-2 as a target whose interaction with TRKB is reduced following acute fluoxetine treatment. In line with a robust interaction between these proteins, we found three out of four subunits of the AP-2 complex interacting with TRKB, while in the case of related complexes, *i.e.* AP-3 and AP-1, only one subunit was co-precipitated for each: AP3D1 and AP1B1. Furthermore, we identified tyrosine-based motifs in the TRKB receptor as putative binding sites for the AP2M subunit of the AP-2 adaptor complex. Especially the region around the Y816 residue of TRKB conformed to the classical consensus for an AP2M-interacting motif. In line with its classical role as endocytic adaptor complex, AP-2 regulates the surface localization of TRKB. Decreased expression of AP2M resulted in increased surface levels of TRKB, facilitating its activation by its ligand BDNF. We also found increased levels of activated TRKB in the hippocampus of AP2M haploinsufficient mice, as well as an increased exposure of TRKB at the surface of cultured hippocampal neurons from these animals. Finally, we observed that classical AD such as fluoxetine, imipramine, rolipram, phenelzine, as well as the fast acting antidepressants ketamine and RR-HNK are able to disrupt the coupling between AP2M and TRKB resulting in increased exposure of TRKB at the cell surface, at concentrations found in the central nervous system after systemic administration (44–46).

It has been established that activation, endocytosis, and trafficking of TRKB receptors regulate the propagation of the downstream signaling response (47). There are three main cascades downstream of TRKB receptor activation: MAPK/ERK signaling regulating neuronal differentiation and growth is activated by recruitment of adaptors to the Y515 site of the receptor. Phosphorylation of the Y515 residue also induces a PI3K/Akt cascade promoting cell survival and growth. Finally, phosphorylation of the Y816 residue is the key step for docking PLC-γ1 which regulates protein kinase C (PKC) isoforms and induces the release of Ca^2+^ from internal sources to activate Ca^+2^/calmodulin (11, 48). Eventually, the post-endocytic sorting machinery plays a major role during the fate-determination process of the receptor signaling (49). In this scenario, some of the vesicles carrying the receptors are destined for retrograde transport towards the cell body (50, 51), while some are recycled back to the cell surface (52). For instance, Pincher, a membrane trafficking protein, regulates the formation of TRK-carrying endosomes from plasma membrane ruffles in soma, axons, and dendrites (53). Pincher-mediated retrograde transport of signaling TRK receptors are rescued from lysosomal degradation (54). Moreover, BDNF-induced survival signaling in hippocampal neurons has been shown to be regulated by Endophilin-A, that is involved in endosomal sorting of active TRKB receptors (39). Evidence suggests that the maintenance of long-term potentiation in hippocampal slices depends on the recycling and re-secretion of BDNF, thus, emphasizing the necessity of TRKB receptor recycling back to the cell surface (55). Another study indicates that Slitrk5, by coupling to TRKB receptors, promotes its recycling to the cell surface in striatal neurons (56).

AP-2, as a member of the endocytic machinery (57), contributes to the formation of clathrin-coated vesicles around cargo proteins for endocytosis (58). In line with this idea, our results indicate a BDNF-induced enhancement of the TRKB:AP2M interaction, suggesting that the receptor is endocytosed upon activation. Although acute treatment with AD (*e.g.* 30 min) can induce a rapid phosphorylation of TRKB receptor in the mouse brain (7), these drugs disrupted the interaction of TRKB:AP2M. Thus, we suggest that the differential regulation of TRKB:AP2M by BDNF and AD may underlie the differential surface expression of the receptor. Zheng and colleagues (27), as well as Sommerfeld *et al* (59), demonstrated that BDNF quickly reduces the TRKB receptor surface level, while our data indicates that antidepressants increase such levels. Interestingly, AMPA receptors, also committed to AP-2 complex-dependent endocytosis in the brain (60, 61), are internalized upon activation by glutamate. However, while the activation of NMDA receptors results in rapid reinsertion of AMPAR subunits to the cell surface, AMPARs activated in the absence of NMDAR activation are destined for degradation in lysosomes (62). Thus, the source of the activation signal recruits alternative machineries to differentially regulate the fate of the target receptor.

Although the AP-2 complex is best known for its major role in endocytosis (63), it remains doubtful whether the main purpose of the identified AP2M:TRKB interaction is facilitated endocytosis of TRKB. A recent study performing endocytosis assays with WT and AP2M null neurons overexpressing EGFP-TRKB and treated with BDNF could not detect any significant defect in the uptake of TRKB upon loss of AP-2. Instead, the authors showed that the retrograde axonal transport of autophagosomes containing TRKB receptor is dependent on AP-2 linking the autophagy adaptor LC3 to p150.Glued, an activator of the dynein motor required for autophagosome transport (19). This led us to conclude that the action of the TRKB-AP-2 complex is not necessarily restricted to the neuronal surface, but can take place in different neuronal compartments. Therefore, antidepressants could disrupt the interaction of TRKB receptors with AP2M at the cell surface, as well as in the autophagosomes.

Disruption of the AP2M:TRKB interaction in internal vesicles could prevent the interaction of the complex with the members of the autophagy-lysosome pathway that can lead to degradation of TRKB receptors (19). The delayed delivery of the signaling endosomes to the soma may locally enhance the specific signaling pathways that can recruit more plasticity-related proteins to the synapses. The two above mentioned scenarios are not exclusive, even though our TRKB:RAB colocalization data points towards changes that take place mainly at the cell surface level. RAB11-positive endosomes have been shown to accumulate in dendrites of hippocampal neurons after BDNF challenge to support dendritic TRKB recycling (40). As we failed to observe an enhancement of TRKB:RAB11 colocalization in spines and neurites under fluoxetine, we suggest that the drug-induced regulation of TRKB surface exposure takes place in the cell membrane, at least shortly after the drug administration. On the other hand, the increase of TRKB:RAB7 colocalization only in spines but not in neurites suggests that TRKB receptors directed towards degradation could be stalled, which could contribute to the maintenance of previously activated and internalized receptors.

In the present study we combined *in vivo, in vitro* and *in silico* methods to gain insight into putative interaction domains between TRKB and AP-2. The *in silico* techniques allowed us to obtain *a priori* knowledge about possible interactions, especially involving the Y816 region and the AP2M subunit. In fact, in the present study, the *in silico* results on the AP2M interaction with the TRKB C-terminus matched the *in vitro* observations. As predicted by the CABS-docking server, AP2M displayed an increased interaction with phosphorylated TRKB (simulated using the phosphomimetic mutation Y816E). Interestingly, the Y816A mutation led to a decrease below the levels of control peptide (WT, non-phosphorylated) observed by both *in silico* (interpreted as a less stable complex) and *in vitro* assays. This suggests that there is some level of interaction between AP2M and non-activated TRKB that is further potentiated upon TRKB phosphorylation. However, the role of such interaction, if any, or its occurrence *in vivo*, remains to be investigated.

At this point, we speculated that the interaction of AP2M and TRKB could be a putative target for AD. Supporting this idea, cell-free assays indicated that the disruption of the TRKB:AP2M complex by fluoxetine happens in a dose-dependent manner. However, the same effect was not replicated in the AP2M:TRKB.peptide interaction assay. Therefore, although our data supports the idea of a binding site for AD in the TRKB complex, it is unlikely that this binding site is between the receptor C-terminal portion and the AP2M subunit. Moreover, this site displays a low affinity for fluoxetine, given that effective doses are found above 0.3 μM. In fact, we have recently identified a low affinity binding site for antidepressants in the transmembrane domain of TRKB (64). However, it is presently unclear how the conformational changes triggered by antidepressant binding to TRKB could alter the interaction of TRKB with AP-2.

Altogether, we suggest a novel mechanism where the antidepressant-induced disruption of the TRKB:AP2M interaction promotes TRKB cell surface exposure and consequently BDNF signaling. However, further investigation will be necessary to address the precise mechanism and the long-term consequences of such a disruption.

## Acknowledgements

The authors would like to thank Sulo Kolehmainen, Seija Lågas (from UH), Claudia Schmidt (from FMP), and the Biomedicum Imaging Unit for their technical assistance; as well as for Dr. Henri Huttunen for RAB5 and RAB7 antibodies. We also thank Drs. Tomi Rantamaki, Cassiano R. A. Faria Diniz, and Caroline Biojone for their helpful comments on the manuscript. Funding for this study was provided by ERC (#322742 – iPLASTICITY), EU Joint Programme - Neurodegenerative Disease Research (JPND) project CircProt (#301225), Academy of Finland (#294710 and #307416), Sigrid Juselius Foundation, Jane and Eljas Erkko Foundation, and Deutsche Forschungsgemeinschaft (DFG SFB958/A01).

## Author contributions

SMF, LL, CB, IC, RM, LV, HG and PCC designed and conducted the experiments. PCC, MV, TM and EC conceived the experimental designs. SMF, LL and PCC wrote the first draft of the manuscript, edited by MV, TM and EC.

